# Using Unmanned Aerial Vehicles (UAV) for free-roaming dog population size estimation

**DOI:** 10.1101/823419

**Authors:** Charlotte Warembourg, Monica Berger-González, Danilo Alvarez, Filipe Maximiano Sousa, Alexis López Hernández, Pablo Roquel, Joe Eyermann, Merlin Benner, Salome Dürr

## Abstract

Population size estimation is performed for several reasons including disease surveillance and control, for example to design adequate control strategies such as vaccination programs or to estimate a vaccination campaign coverage. In this study, we aimed at assessing the benefits and challenges of using Unmanned Aerial Vehicles (UAV) to estimate the size of free-roaming domestic dog (FRDD) populations and compare the results with two regularly used methods for population estimations: a Bayesian statistical model based on capture-recapture data and the human:dog ratio estimation. Three studies sites of one square kilometer were selected in Petén department, Guatemala. UAV flight were conducted twice during two consecutive days per study site. The UAV’s camera was set to regularly take pictures and cover the entire surface of the selected areas. A door-to-door survey was conducted in the same areas, all available dogs were marked with a collar and owner were interviewed. Simultaneously to the UAV’s flight, transect walks were performed and the number of collared and non-collared dogs were recorded. Data collected during the interviews and the number of dogs counted during the transect walks informed a Bayesian statistical model. The number of dogs counted on the UAV’s pictures and the estimates given by the Bayesian statistical model, as well as the estimates derived from using a 5:1 human:dog ratio were compared to dog census data. FRDD could be detected using the UAV’s method. However, the method lacked of sensitivity, which could be overcome by choosing the flight timing and the study area wisely, or using infrared camera or automatic detection of the dogs. We also suggest to combine UAV and capture-recapture methods to obtain reliable FRDD population size estimated. This publication may provide helpful directions to design dog population size estimation methods using UAV.

## Introduction

Estimating the size of domestic and wild animal populations has been used in diverse research fields, such as for species monitoring and conservation [1,2] or disease surveillance and control [3]. For example, population size estimation methods have been used to monitor biological diversity in space and time [4], estimate the abundance and conservation status of rare and endangered species such as whales or rare and elusive carnivores [5,6] or to design and conduct vaccination programs [7,8] and estimate vaccination campaign coverage for infectious diseases [9].

Methods have been developed to estimate the size of both wild [10] and domestic animal populations [11,12]. Among domestic animals, stray dog populations have been widely studied because of their importance in the transmission of rabies, a deadly zoonotic disease [13–17]. According to the World Health Organization (WHO), more than 99% of human rabies cases worldwide are caused by dog bites [18]. The World Organization for Animal Health (OIE) defines stray dogs as dogs not under direct control by humans or not prevented from roaming freely. They are divided into three categories: free-roaming owned dogs, free-roaming ownerless dogs still living within human communities and feral dogs which have reverted to a wild state and do not depend on humans anymore [19]. The dogs of the first two categories are also named as “free-roaming” or “free-ranging”, which encompass both owned and ownerless dogs [20] and are more relevant for zoonotic disease control than feral dogs due to their closeness to the human population.

The estimation of free-roaming domestic dog (FRDD) population size is challenging and susceptible to bias because of the presence of heterogeneity in the dog population (i.e. presence of owned and ownerless dogs) which leads to heterogeneity in detection probabilities [13]. The most commonly used methods are based on census surveys [21,22], distance sampling (transect based method where perpendicular distances from random transect lines to the dogs are measured to estimate the dog density) [14,23] and capture-recapture technics [14–17,20,24]. However, most of these methods do not account for the presence of ownerless dogs and may therefore underestimate the total dog population size to be considered for disease control, such as rabies. Other methods, for example Bayesian statistical models, are able to account for heterogeneity of the population by considering different detection probabilities for subpopulations and the use of prior information on the proportion of ownerless dogs [9,25].

Guatemala is a Central American country where canine rabies is still endemic. Ministry of Health reported 956 animal rabies cases and 18 human rabies deaths in the country during the period of 2005-2017 [26]. The current dog population size in Guatemala is unknown. A study conducted in 2008, focusing on owned dogs, estimated the number of dogs in 12 neighborhoods of Todos Santos Cuchumatán, a town located in the Department of Huehuetenango, at 392 [27]. The method chosen was a door-to-door household census. Based on a study published in 1959 [28], the animal health authorities in Guatemala currently use a human:dog ratio of 5:1 to estimate the number of vaccines needed for the national rabies vaccination campaigns. However, the human:dog ratio may vary regionally, which was also found as preliminary outcome of currently ongoing studies undertaken by the Universidad del Valle (UVG) that aim to estimate the human:dog ratio in urban, peri-urban and rural communities (unpublished data). The methods used so far to estimate the dog population size in Guatemala focused on owned dogs only and did not take into account the potential presence of ownerless dogs within the population.

Thanks to recent progress in technology and software development, the utilization of Unmanned Aerial Vehicles (UAV) that can take geo-referenced pictures and automatically fly along pre-defined paths has strongly increased for population monitoring [29,30]. Commonly called drones, they became a valuable tool in agricultural, environmental and wildlife monitoring [31]. Some studies used UAV to estimate population sizes of various animal species including sea birds [29,32], marine fauna [29,33], biungulates (red deer, roe deer, elk, wild boars, bison) [34], wolves [35] and marsupials [31]. Comparing to other counting methods, UAV’s advantage is the ability to count wildlife time efficiently in remote areas and in a standardized and reproducible way [35,36]. These features may be of interest for FRDD counting, especially for dogs that do not have an owner and might have an elusive behavior with human beings. To the best of our knowledge, this is the first study published that evaluates the possibility of using UAV to estimate the size of FRDD populations.

The aim of this study is to assess the benefits and challenges of using UAV to estimate the size of FRDD populations in three locations in Guatemala and compare the resulting estimates with those of two regularly used methods for population estimations. These methods are a capture recapture technique using Bayesian statistical model and the utilization of the human:dog ratio. Practicability and ethical considerations of the use of drones will be discussed as well.

## Material and methods

### Study area

The study was performed in the Petén Department of Guatemala, located in the Northern lowlands of the country (Fig 1), specifically in the Poptún Municipality, one of 14 political subregions. It covers approximately 69,000 square kilometers (31.8% of Guatemala’s surface) and constitutes the southern part of the Yucatan tableland [37]. It is a sub-tropical forest biome with thick carbonate deposits responsible for a predominant karst topography, creating thin soils that erode easily and are not fit for agriculture [38]. Once scarcely populated, this region has seen mass migration patterns increase in the last 30 years, causing an accelerated land-use change associated to large scale African palm production, cattle industry and slash-and-burn agriculture [39]. Population in the region is 69% Ladino (Spanish-speaking) and 30% Maya, with an overall illiteracy rate of 22%. Close to half of the population is living under the poverty line [40]. Most of the populated areas are considered rural (59.46%), where access to basic services such as safe drinking water, electricity, sanitation, public health facilities, or transportation is limited [40].

**Fig 1:**
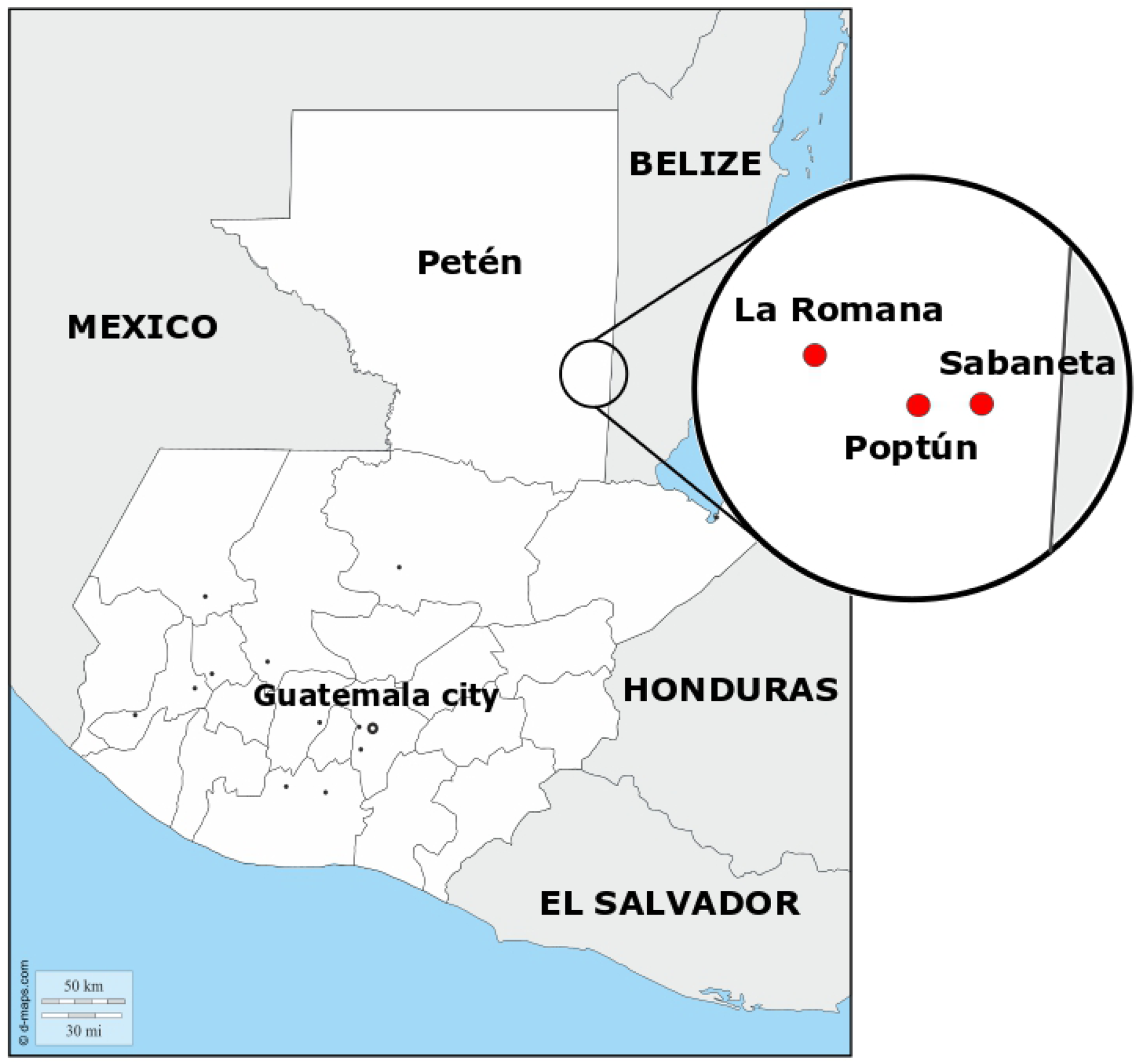
Localization of the three study areas (Poptún, Sabaneta and La Romana) in Petén department, Guatemala. Source: d-maps.com

Three study areas were selected within the Poptún Municipality, depending on their expected dog population sizes. Overall, Poptún has 52,282 inhabitants according to the 2018 census (INE, 2019). The main city of Poptún, categorized as an urban area, has an estimated population of 29,000 people. Sabaneta and La Romana are two villages north of Poptún with 804 and 485 residents, respectively, surrounded by agricultural landscape (rural areas). Sabaneta and La Romana were selected by convenience as they were already involved in a research project named “One Health Poptún” implemented by researchers of the UVG(http://www.ces.uvg.edu.gt/page/2018/02/07/proyecto-una-salud-poptun-video/). This project aimed to improve the health situation of Maya communities in the Poptún Municipality using a One Health approach. Poptún city was chosen as it was the closest urban area to Sabaneta and La Romana.

As part of “One Health Poptún” research project, a human and domestic animal census was performed in Sabaneta between June 2017 and April 2018 and in La Romana between June 2017 and July 2018. A total of 110 and 289 owned dogs were recorded in La Romana and Sabaneta, respectively. No dog census data was available for Poptún.

For each of the three study areas, we defined a one square kilometer study site, in which the dog population estimation was performed. In La Romana and Sabaneta, the study sites covered the entire village; in Poptún, it only covered part of the city. As study population, we considered all type of dogs visible in the streets or yards regardless of their ownership status. It therefore constituted of owned or ownerless FRDD or stray dogs as described in the OIE guidelines on stray dog population [19]. The study was conducted between May and June 2018.

### Ethical approval

Given the study design included research with human subjects as well as with animals, the research team submitted two ethical protocols prior to conducting fieldwork, one to UVG’s International Animal Care and Use Committee (IACUC), and another to the Ethics Review Board (ERB) of the Committee for Research on Human Subjects of the Center for Health Studies in UVG. Since the methodology included the use of UAV to fly over both private and public areas in each town, it was necessary to obtain permits from the National Authority for Civil Aeronautics (DGAC in Spanish), which included registering each drone with a unique identification number. Given research with UAVs is relatively new in Guatemala, the DGAC had no written regulations pertaining to ethical considerations for their use in contexts like the one addressed by this project, a void commonly reported in the scarce literature on UAVs for research [41]. The lack of clear guidelines to address issues of privacy, data protection and public safety, along with conducting research in indigenous populations considered vulnerable [42], demanded this project followed consuetudinary law as practiced in each site. The research design was presented and discussed with health and political authorities of Poptún in several meetings until all concerns for public safety were addressed. Official research permits were then requested and obtained from the Regional Health Authority (Petén Sur Oriente) of the Guatemala Ministry of Health and from the Municipal Government of Poptún. The latter granted specific permits to conduct research in the urban area and directed the communication campaign to inform citizens about the UAV flights, selection of homes and related activities. While the urban area is ruled only by municipal law and has a predominant Ladino population, the towns of Sabaneta and La Romana are predominantly Maya Q’eqchi’ (65% and 98% respectively) and are ruled by traditional indigenous consuetudinary law. In both sites, the team met with local leaders from the Community Development Councils (COCODES) during March and April 2018 to explain the aims and methods of the research, address concerns, and request formal authorization for implementation. In La Romana, the COCODE asked that the team returns on a following weekend to explain to the entire population the research endeavor in an Open Council, in the local language, where a voting and consensus process was conducted resulting in the community’s approval of the project. This decision was formally communicated to the research team one week later in writing by the COCODE leaders. In Sabaneta, the COCODE authorities determined the intervention was of value to the community, but requested a door-to-door campaign to explain to each family head the proposed research. Once this process was concluded, the COCODE extended a formal letter of approval to conduct the research. The process was led by anthropologists and supported by Maya Q’eqchi’ staff of UVG to guarantee cultural pertinence, enhance two-way communication, build trust, and understand compliance to consuetudinary law. Local community authorities were involved in on-site supervision of all research activities.

### Dog population size estimation

Three methods were used to estimate the dog population sizes in the three study areas and results were compared.

#### Method 1: Utilization of UAV

The UAV used in this study was a DJI Mavic Pro (https://www.dji.com/ch/mavic). The main characteristics of its aircraft and camera are described in Table 1.

**Table 1:**
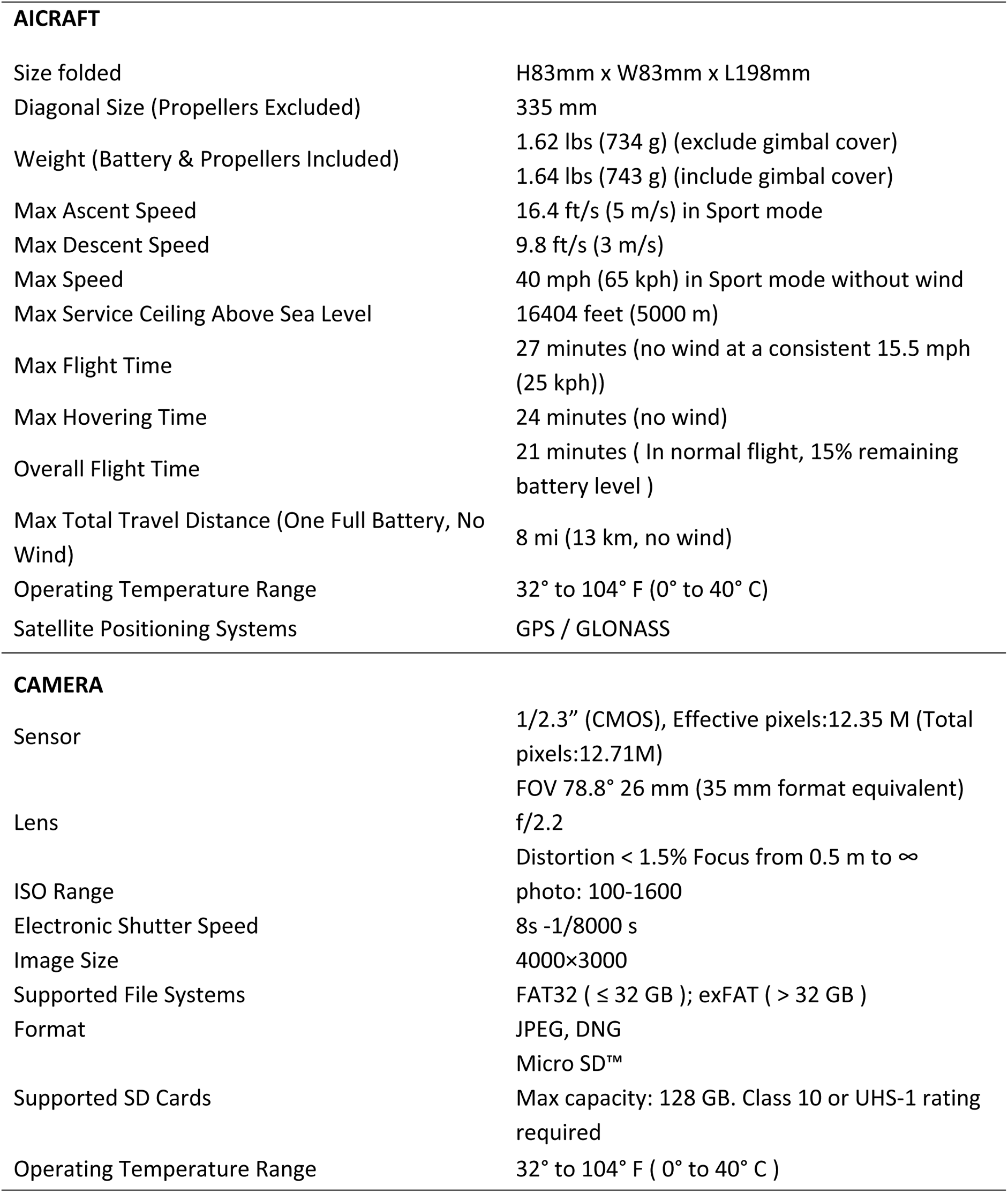
Technical specifies of the Mavic Pro used for the UAV flights.

A total of twelve flights were scheduled, four per study site. In each site, the flights were conducted twice a day, in the morning and late afternoon, over two consecutive days. The flight course was mapped in advance using Pix4Dcapture, a free mobile application for drone flight planning (https://www.pix4d.com/product/pix4dcapture). A grid with parallel transect lines covering the predefined flight area were defined automatically by the application. The number of transect lines was conditioned by the altitude of the flight and pictures’ overlap selected by the operator. Accordingly, the number of pictures is given by the altitude of the flight and pictures’ overlap, ensuring that the entire study site surface is captured by the pictures. Those parameters were dependent on the number and capacity of batteries available. Five batteries (capacity: 3038 mAh, voltage: 11.4V, battery type: LiPo 3S, Net weight: approx. 240g) were available per flight.

Table 2 shows the overlap, altitude and number of pictures taken per flight in the three study areas. The pictures were automatically stored in the UAV’s SD card.

**Table 2.**
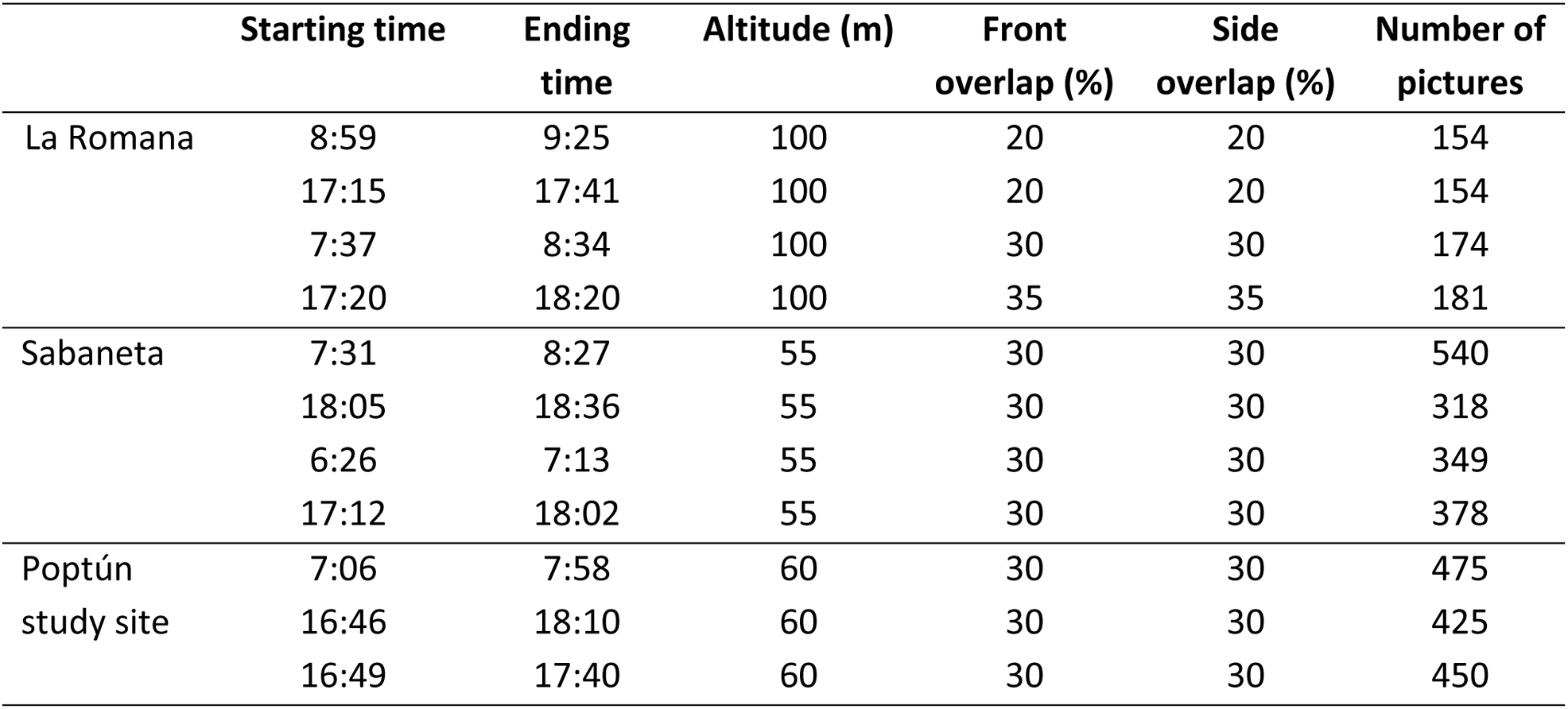
Characteristics of the UAV flights performed in Petén department, Guatemala in May-June 2018. Front and side overlap are defined by the overlap area of two consecutive pictures along the flight line (front) and between transect lines (side). In Poptún, only three flight could be realized because of bad weather conditions during the last flight.

|  | Starting time | Ending time | Altitude (m) | Front overlap (%) | Side overlap (%) | Number of pictures |
| --- | --- | --- | --- | --- | --- | --- |
| La Romana | 8:59 | 9:25 | 100 | 20 | 20 | 154 |
|  | 17:15 | 17:41 | 100 | 20 | 20 | 154 |
|  | 7:37 | 8:34 | 100 | 30 | 30 | 174 |
|  | 17:20 | 18:20 | 100 | 35 | 35 | 181 |
| Sabaneta | 7:31 | 8:27 | 55 | 30 | 30 | 540 |
|  | 18:05 | 18:36 | 55 | 30 | 30 | 318 |
|  | 6:26 | 7:13 | 55 | 30 | 30 | 349 |
|  | 17:12 | 18:02 | 55 | 30 | 30 | 378 |
| Poptún study site | 7:06 | 7:58 | 60 | 30 | 30 | 475 |
|  | 16:46 | 18:10 | 60 | 30 | 30 | 425 |
|  | 16:49 | 17:40 | 60 | 30 | 30 | 450 |

In each study site, a team of two persons conducted the UAV flights, the operator and the observer. The operator was in charge of the controller and monitored the flight on the control screen. The camera view was displayed on the control screen and a signal was transmitted each time a picture was taken. The operator checked that the transmission between the UAV and the controller was functioning well and pictures were regularly taken. There is no need to drive the UAV as it flies automatically following the flight course defined on Pix4Dcapture application. The observer was standing next to the operator. He was visually controlling that the UAV was regularly visible, operating well and did not enter in collision with a bird, a tree or an antenna. When the UAV ran out of battery, it automatically returned to the take-off spot and the observer replaced the battery. Then, the UAV flew back to the previous location and pursued with the data collection.

The analysis of the pictures was undertaken after the data collection period by researchers from the UVG Arbovirus and Zoonosis group. The first step was to eliminate pictures that could not be analysed due to poor quality (blurred or taken at too high altitude) or fully covered by forest canopy. This exclusion phase was done by three people simultaneously. The rest of the pictures was analysed individually to identify dogs. When there was any doubt about the identification of a dog in any of the pictures, these were analysed again by three different people to have a consensus on whether there was a dog or not. The pictures to be further reviewed were then stored electronically sorted by study area, date and time.

To identify dogs including their location, each picture was divided into nine (3 x 3) quadrants and each quadrant was identified with a combination of letters and numbers (Fig 2). Each picture and quadrant was reviewed in a zigzag pattern, from top left to bottom right, and identified dogs were marked with a red circle. Doubtful observations resembling dogs but not presenting dog characteristics, such as ears, thorax, pelvis, limbs or tails, were not counted as dogs.

**Fig 2:**
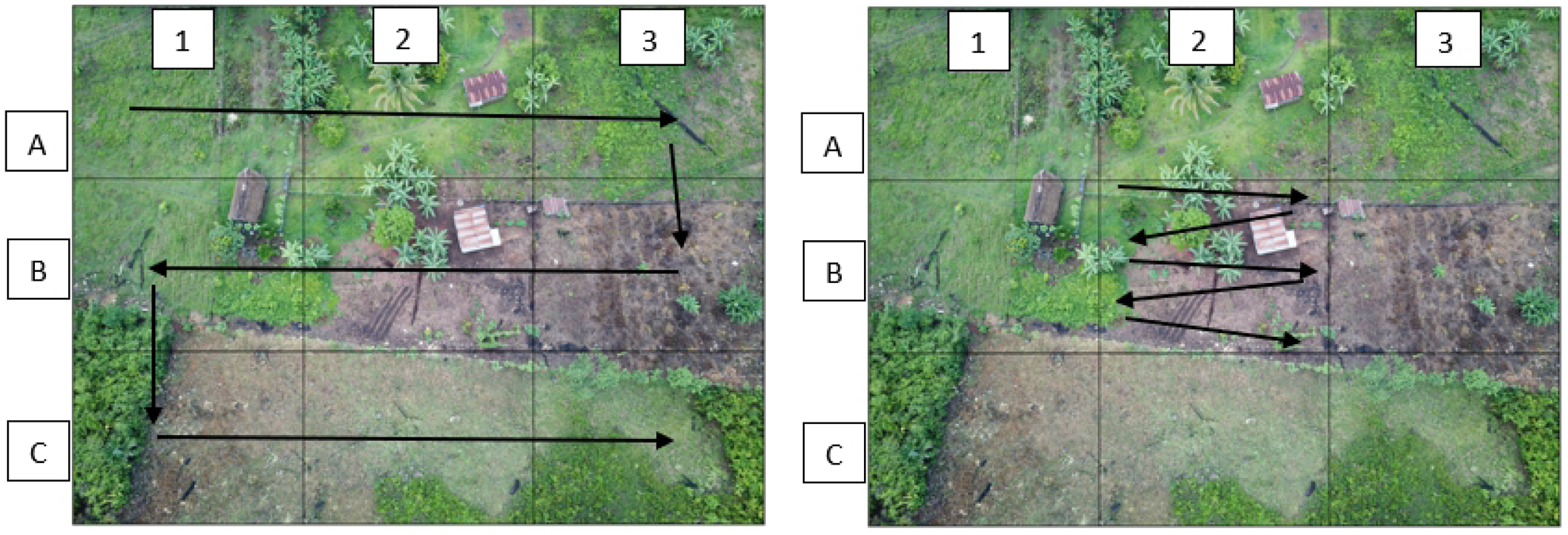
Reviewing procedure of picture taken by the UAV in one of the rural sites. The left picture shows the nine quadrants and the zigzag pattern used to review the picture. The right picture shows the zigzag pattern used to review each quadrant.

The study area, identification number of the picture, date and time when the picture was taken, identification of the dog, location of the dog in the quadrant, general observations on the dog (e.g. fur colour, presence of a collar), reviewer’s name and date of review were entered in an Excel document for each identified dog.

#### Method 2: Bayesian statistical model based on capture-recapture data

Data required for the Bayesian statistical model include capture-recapture data, number of collar lost during the study, prior information on the dog confinement, on the ratio of owned versus ownerless dogs and on the number of owned dogs in the study sites. These data were collected during transects (capture-recapture data) and a household survey (dog confinement, proportion of ownerless dogs).

The week before the UAV’s flights, all households located within the three study sites were visited and dog-owning households were identified. In each dog-owning household, we interviewed the dog owner and aimed to mark all dogs of the household by a collar. The collars used for marking the dogs were red and carried a grey box containing a GPS chip (used for another study, no meaning here), easily visible from several dozen meters. In Sabaneta and La Romana, we collared both free-roaming dogs and confined dogs if they could be in contact with FRDD dogs (e.g. restrained by a chain in an open yard). In Poptún, due to the high number of dogs, the team focused the collaring on free-roaming dogs and not on fully confined dogs. The reasons for non-collaring of the dogs were refusal of the owner to participate to the study, dog’s stress or aggressiveness, or absence of the owner or the dog during the visit. A small proportion of the households refused to participate and the reasons for not participating included cultural beliefs and owners thinking that the collar would disturb the dog during hunting. Dogs younger than four months, pregnant bitches and sick animals were excluded from the study and not marked. The reasons for this was the too small size of the puppies for the collars and to avoid unnecessary stress of pregnant or sick animals.

The interviews were conducted using a questionnaire collecting information on dog confinement practices and owner estimates on the number of owned dog, community dog and ownerless dog populations in their living area. A dog with a dedicated owner is considered as an owned dog, a dog without owner but fed by the community is considered as a community dog and a dog without owner and not fed by the community or a particular person is considered as an ownerless dog. The data were directly put in a digital format into a tablet equipped with KoBoCollect Android application (https://www.kobotoolbox.org) and later downloaded as an Excel spreadsheet. In total, 61 dogs were marked in La Romana, 125 in Sabaneta and 117 in Poptún study site, within 39, 76 and 73 households respectively, which corresponds to the number of interviews conducted. The data collection lasted for four days in Poptún and Sabaneta study sites and three days in La Romana.

Simultaneously to the UAV flights, a second team performed ground transect walks to count dogs in the streets and opened yards where dogs had access to the street. The team was constituted of at least three persons: a local guide, a person recording the number of dogs and a person spotting the dogs. The transect lines were defined in advance on Google Earth according to the following rules (Figs 1-3 - supporting information). The minimum distance between two lines was 100 meters, a transect line should not cross another line and we respected a buffer zone at the edge of the site of 100 to 400 meters to avoid counting dogs whose owner is not living in the study site. While counting the dogs, we distinguished between marked (collared dogs spotted during the transect walks) and non-marked dogs (dogs spotted during the transect walks not carrying the study collars). To avoid double counting of individual dogs, we took pictures or videos of the dogs encountered. During the transects, if the team had a doubt whether a dog was being seen for the second time, they would go through the pictures taken previously, check if the dog had already been counted and decide whether they count or ignore it. The total transect length was 3 kilometers in La Romana, 4.6 kilometers in Sabaneta and 8.2 kilometer in Poptún study sites and the mean time for conducting the transect was 58 minutes, 71 minutes and 128 minutes, respectively.

**Fig 3.**
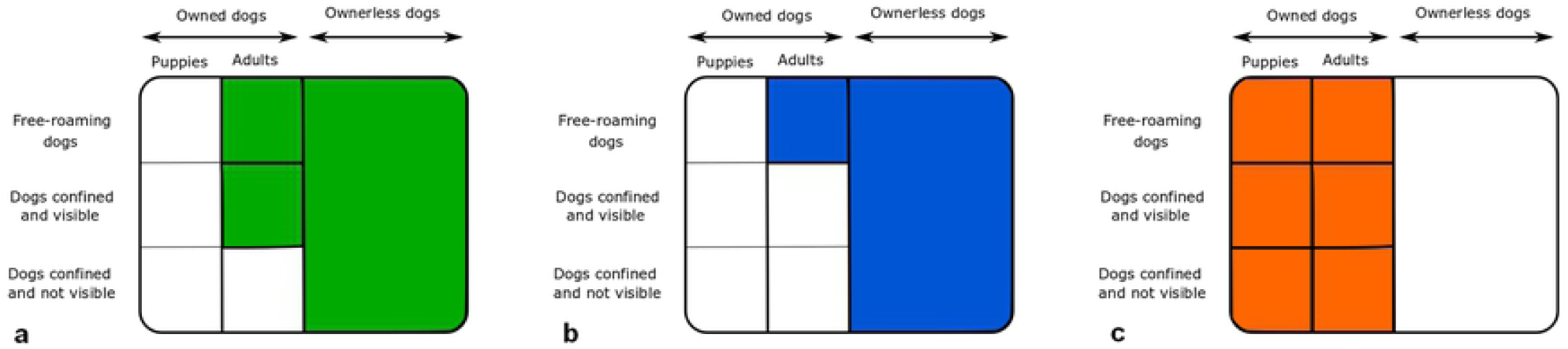
Structure of the dog population in the study region, Petén department, Guatemala.

A Bayesian statistical model (File 1 – supporting information) was developed to estimate the dog population size based on number of dogs marked during the household study, number of dogs spotted during the transect walks (distinguishing between marked and unmarked dogs), the number of collars lost during the study and study site specific prior information derived from the questionnaire data. The model considers the probability of a dog to be spotted during the transect walks, the probability for a marked dog to loose the mark (i.e. collar), the probability of a dog being confined (and therefore not seen) at the time of the transect walks and the probability of encountering ownerless dogs during the transects.

All the marked dogs encountered during the transect walks were owned, whereas the unmarked dogs either were owned (but unmarked) or ownerless. Since we assumed that we marked more than 10% of the dog population, we considered that the number of marked dogs, owned unmarked dogs and ownerless unmarked dogs counted during the transects follow a hypergeometric distribution [43]. Probability mass function p(x) of the hypergeometric distribution model with n denoting the total number of dogs spotted during the transect walks, D the number of dogs marked during the study, M the total size of the dog population and x the number of marked dogs encountered during the transect walks as following [44]:

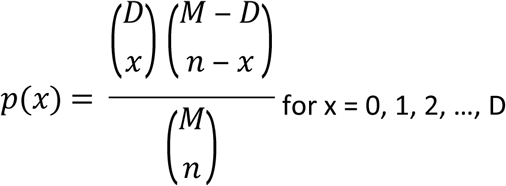

where 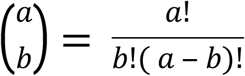

The probability lm(t) that a marked dog lost its collar at transect number t was modelled as:

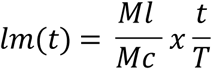

with Ml denoting the total number of collars lost during the entire study period (three in La Romana, two in Sabaneta, four in Poptún study site); Mc denoting the total number of dogs collared in the study (61 in La Romana, 125 in Sabaneta and 117 in Poptún study site) and T denoting the total number of transects per study site (four each in La Romana, Sabaneta and Poptún).

We considered that ownerless dogs have a confinement probability of 0. Prior data on the confinement probability of owned dogs were calculated based in the information collected during the interview survey. The dog owner was asked to explain when their dog usually roams. The answer was then categorized into “never free-roaming”, “always free-roaming”, “free-roaming during the day only”, “free-roaming during the night only”, “free-roaming a few hours per day only” (Table 1 - supporting information). We considered confinement probabilities during the time of the transects of 1 for dogs of the category “never free-roaming”, 0 for dogs of the category “always free-roaming”, 0-0.25 (uniform distribution) for dogs of the category “free-roaming during the day only”, 0.5 for dogs of the category “free-roaming a few hours per day”, and 0.75-1 (uniform distribution) for dogs of the category “free roaming during the night only”. The minimum and maximum confinement probability were calculated for each study area based on these probabilities, considering the lower and upper limit of the range, respectively, weighted by the number of dogs per category. The prior information of the dog confinement probability was included into the Bayesian model as a study site specific uniform distribution ranging from the minimum to the maximum confinement probability (Table 3).

**Table 3.** Prior information used in the Bayesian statistical model.

|  | La Romana | Sabaneta | Poptún |
| --- | --- | --- | --- |
| Confinement probability (uniform distribution) |  |  |  |
| Min | 0.133 | 0.157 | 0.293 |
| Max | 0.142 | 0.157 | 0.320 |
| Ownerless to owned dog ratio (log normal distribution) |  |  |  |
| $\mu$ | -6.57 | -7.71 | -7.67 |
| $\tau$ | 5.24 | 4.97 | 4.98 |
| Number of owned dogs living in the study site (uniform distribution) |  |  |  |
| Min | 69 | 175 | 172 |
| Max | 1000 | 1000 | 1000 |

Prior data on the probability of encountering an ownerless dog during the transects was calculated based on the owners’ estimates on the number of owned and ownerless dogs living in the study sites requested during the interview survey. We then calculated, for each study area, the ratio of ownerless to owned dogs using the estimation given by the respondents (Fig 4 – supporting information). Log normal distributions were fitted on the data and distribution parameters were used as prior information incorporated in the statistical model (Table 3).

**Fig 4.**
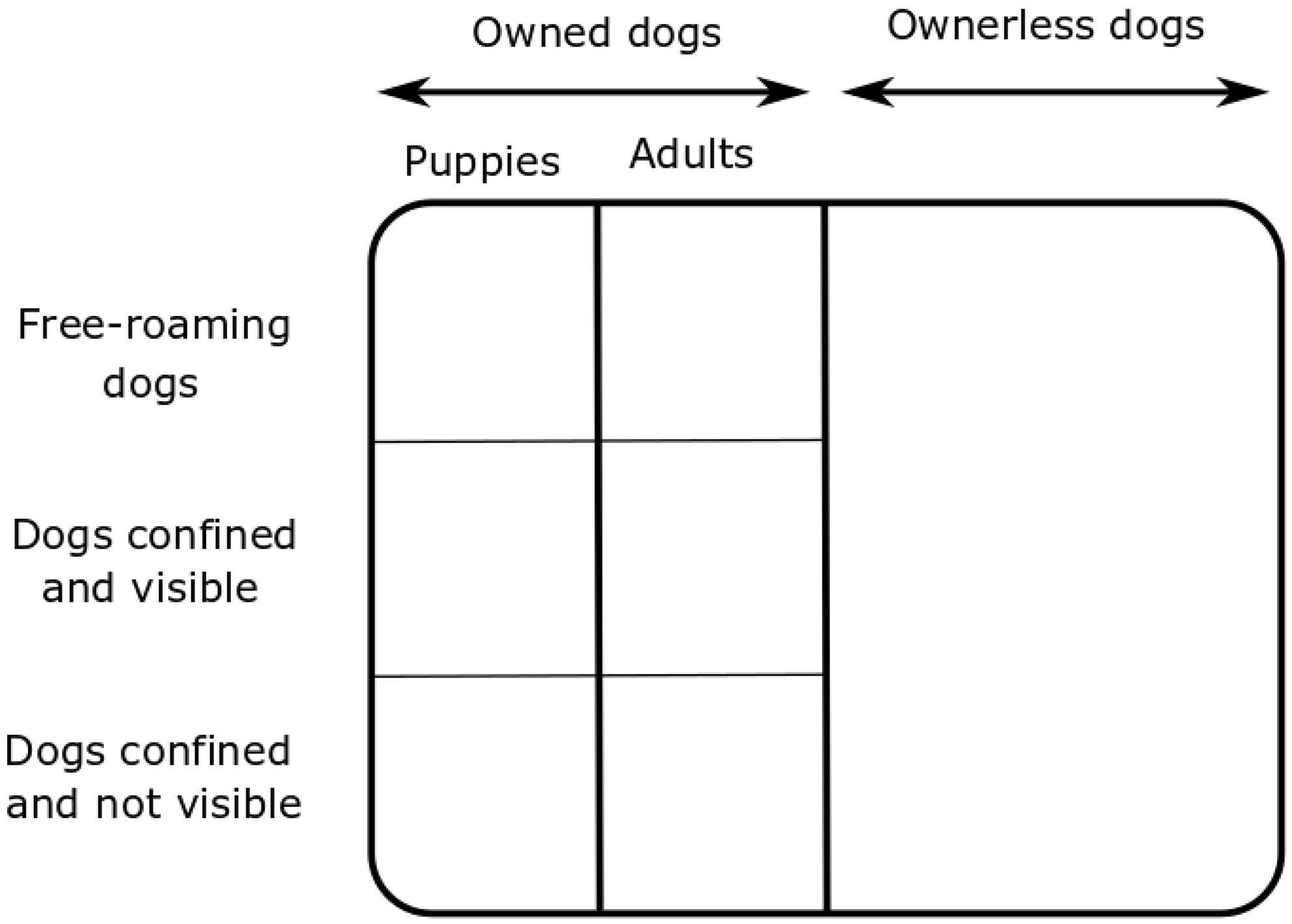
Study population of the methods used to estimate the size of dog populations. a. Method using UAV, b. capture-recapture method and analysis using Bayesian statistical model, c. dog census and human:dog ratio method.

No prior data was collected on the number of owned dogs during the interviews. Therefore, we intended to use non-informative priors. The number of owned dogs was modeled as a uniform distribution. The maximum value was set up at 1,000 in each study area, which was considerably higher than any expectations. A theoretical minimum value was considered as the number of dogs collared during the study in each study area. However, this number was too low in the three study areas to initiate the Bayesian model. Therefore, the lowest number that was practically possible for model initiation was used in the three study areas (Table 3).

The model was implemented in OpenBugs (http://www.openbugs.net) version 3.2.3. The code of the model is presented in the supporting information.

#### Method 3: the human:dog ratio

Applying the human:dog ratio on the human population size in each study area builds the third method to estimate the dog population size. We used the human:dog ratio currently used by the health authorities in Guatemala for the annual national rabies vaccination campaign, 5:1 [28].The number of humans was derived from the total human census performed by the project “One Health Poptún” in Sabaneta and La Romana in 2017-2018, leading to a population size of 804 and 485 people, respectively. Since, the study areas in La Romana and Sabaneta cover the entire villages, we used the total population to calculate the number of dogs. Because of lack of human population data in Poptún study site, we could not estimate the size of the dog population in Poptún study area using the human:dog ratio method.

## Results

Out of the twelve flights, only eleven could be realized because of bad weather conditions (rain) during the third UAV flight in Poptún. A total of 3,407 pictures were reviewed from the 11 flights: 520 in La Romana, 1,537 in Sabaneta, and 1,350 in Poptún study site. Out of them, 2,685 were selected for review, 293 from La Romana, 1,048 from Sabaneta and 1,344 from Poptún. In the pictures, it was easier to detect dogs located on the roads than in household patios, in the shade, under a tree or under other constructions. When dogs were located in closed yards, it was not possible to differentiate partly free-roaming dogs and completely confined dogs. Because of their small size and their tendency to stay at home before weaning, young puppies were not spotted on the UAV’s pictures. The number of dogs countered per flight is shown in Table 4. The numbers of dogs encountered during the transects walks and used for the Bayesian statistical model are also presented for comparison.

**Table 4.**
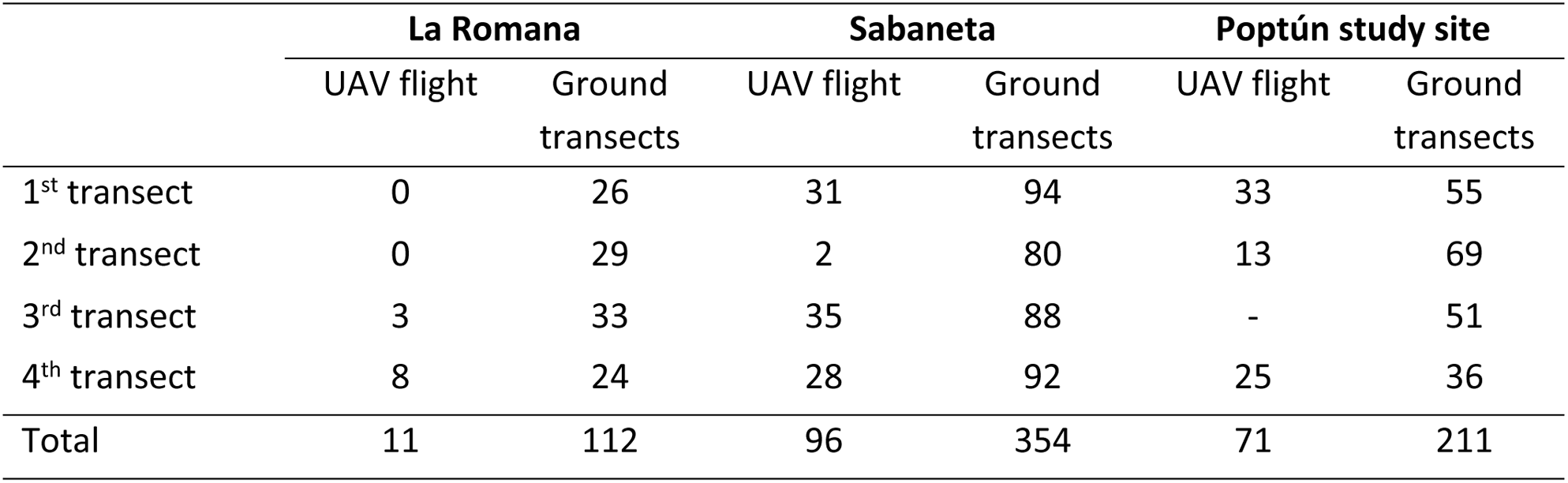
Number of dogs counted during UAV flights in comparison with the number of dogs spotted during the ground transect walks in three study areas of Poptún Municipality, Petén department, Guatemala in May-June 2018.

|  | La Romana |  | Sabaneta |  | Poptún study site |  |
| --- | --- | --- | --- | --- | --- | --- |
|  | UAV flight | Ground transects | UAV flight | Ground transects | UAV flight | Ground transects |
| 1<sup>st</sup> transect | 0 | 26 | 31 | 94 | 33 | 55 |
| 2<sup>nd</sup> transect | 0 | 29 | 2 | 80 | 13 | 69 |
| 3<sup>rd</sup> transect | 3 | 33 | 35 | 88 | - | 51 |
| 4<sup>th</sup> transect | 8 | 24 | 28 | 92 | 25 | 36 |
| Total | 11 | 112 | 96 | 354 | 71 | 211 |

Substantially more dogs were observed during the transect walks than on the pictures taken by the UAV. This difference is not caused by counting puppies during the transects, because spotting young puppies was very unlikely due to their tendency to stay inside with the bitch and were not be visible from outside of the compounds.

The Bayesian model estimated a total dog population size of 74.7 in La Romana, 262.6 in Sabaneta and 500.8 in Poptún study sites. The credibility intervals and the estimated number of owned and ownerless dogs are shown in Table 5. The number of marked and unmarked dogs recaptured during the transect walks is shown in Table 2 of the supporting information.

**Table 5:**
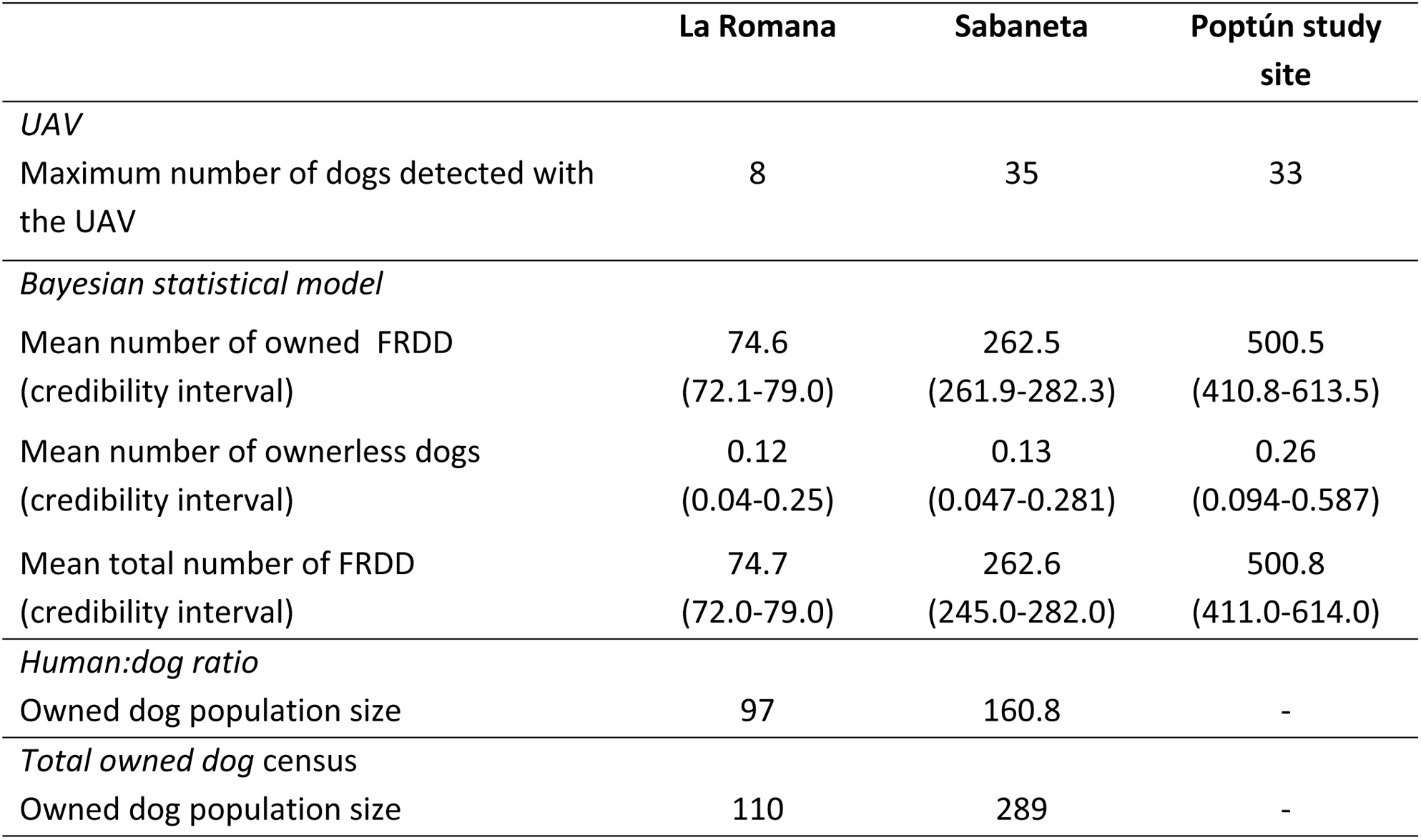
Dog population size estimation given by the UAV, the Bayesian statistical model, the human:dog ratio and the total owned dog census collected during the “One Health Poptún” project.

|  | La Romana | Sabaneta | Poptún study site |
| --- | --- | --- | --- |
| *UAV* |  |  |  |
| Maximum number of dogs detected with the UAV | 8 | 35 | 33 |
| *Bayesian statistical model* |  |  |  |
| Mean number of owned FRDD (credibility interval) | 74.6<br>(72.1-79.0) | 262.5<br>(261.9-282.3) | 500.5<br>(410.8-613.5) |
| Mean number of ownerless dogs (credibility interval) | 0.12<br>(0.04-0.25) | 0.13<br>(0.047-0.281) | 0.26<br>(0.094-0.587) |
| Mean total number of FRDD (credibility interval) | 74.7<br>(72.0-79.0) | 262.6<br>(245.0-282.0) | 500.8<br>(411.0-614.0) |
| *Human:dog ratio* |  |  |  |
| Owned dog population size | 97 | 160.8 | - |
| *Total owned dog census* |  |  |  |
| Owned dog population size | 110 | 289 | - |

Using the human:dog ratio, we obtained an estimation of 97 dogs in La Romana and 160.8 dogs in Sabaneta (Table 5). These numbers were compared to the census data of owned dogs collected during the “One Health Poptún” project.

## Discussion

In this study, three methods estimating dog population size in three study areas in Guatemala were compared. This is the first published study that applied UAV to estimate population size for this species.

To correctly interpret the comparison of the results of the three methods, we need to consider the heterogeneity of the dog population and which subpopulation each method is quantifying. The total dog population can be divided into owned and ownerless dogs (Fig 3). Further, the owned dogs are free-roaming (partly or completely), completely confined and visible (e.g. in a yard fenced with a grid) or completely confined and not visible at any time (e.g. under a roof inside a compound or kept in the house). The owned dogs further have to be separated into puppies (less than 3 months) and the adult dogs.

The UAV are able to spot ownerless dogs, owned free-roaming dogs and owned confined visible dogs but not puppies, which are too small in size (Fig 4a). The Bayesian model is designed to count ownerless and owned free-roaming dogs (partially or completely) only and therefore do not consider confined dogs (Fig 4b). Puppies less than 4 months old were not considered in the Bayesian statistical method because they could not be marked (too small in size for the collar) nor spotted during the transect walks (not visible because they tend to stay at home with the bitch). Unlike the two previous methods, the dog census and human: dog ratio methods focus on all types of owned dogs, including puppies, and do not consider ownerless dogs (Fig 4c).

The human:dog ratio method and the dog census detect the same dog populations, thus the results can be directly compared. In our study, the human:dog ratio estimated a population size close to the census value (97/110 = 88%) in La Romana and underestimated the owned dog population size in Sabaneta, taking the census data as a reference (161/289 = 56%). The Bayesian statistical model performed better than the method using UAV with respectively 68% (75/110) and 7% (8/110) of the dogs detected in Sabaneta and 91% (262/289) and 12% (35/289) in La Romana, taking the census data as a reference. These two methods almost cover the same study population, considering the small proportion of dogs confined (Table 1 – Supporting information) and the estimated negligible number of ownerless dogs (Table 5) in La Romana and Sabaneta (Fig 5). We therefore highlight that when comparing FRDD population size estimates to first carefully evaluate the subpopulation captured by the respective method used.

**Fig 5.**
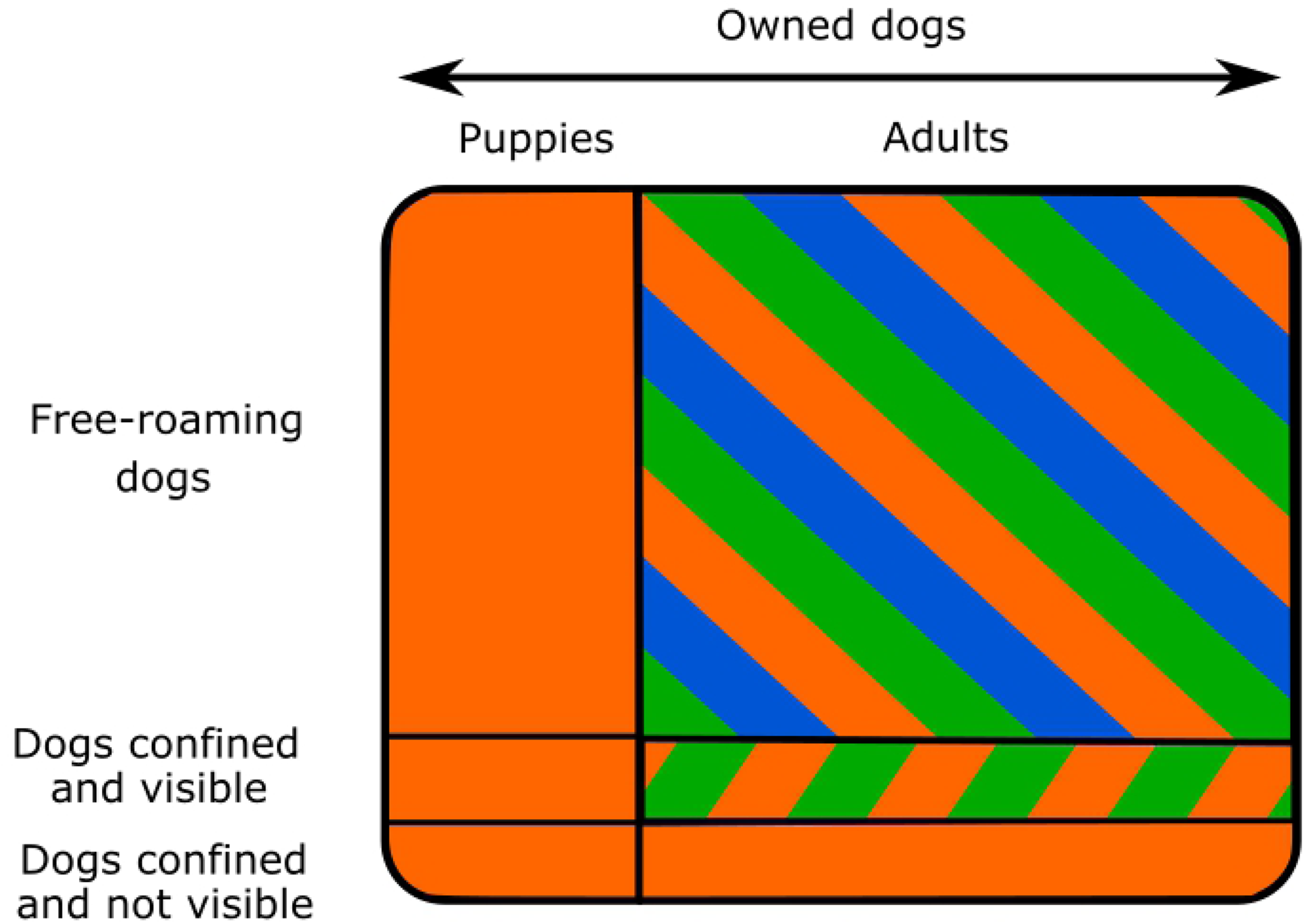
Structure of the dog population in La Romana and Sabaneta with approximate proportions of the sizes of the subpopulations. The ownerless dogs are negligible. The subpopulation captured by the respective method is represented in orange for the dog census, green for the method using UAV and blue for the capture-recapture method.

### Method 1: Utilization of UAV

In this study, we were able to demonstrate that detecting and counting FRDD using UAV is possible. However, we observed a very low sensitivity of this method and could only observe a maximum of 12% of the dogs counted during the dog census. The suggested main reasons for this large underestimation are: a) dogs are not visible from the air by the UAV at the time of the flights, b) the altitude of the flight is too high to identify the dogs and c) the quality of the pictures is poor. In the following section, we will develop these three limitations (a-c) and describe potential solutions.

The dogs were not visible by the UAV at the time of transects when they were located under a tree, a roof overhang or any structure that covers the dog. Dogs tend to hide in cooler place when the weather is hot. To overcome this problem, three potential solutions can be listed: optimizing selection of study site and time of flights, use infrared thermal imagery and combine UAV with capture-recaptured methods.

To increase the proportion of dogs detected in the visible and the infrared spectrum, UAV flights should be performed in a fully open environment and when the dogs are more likely to be roaming, early morning and late afternoon [45]. Local community leaders later suggested flying the drones at sunrise, when dogs look for the first rays of sunlight to warm up from the cold night and therefore leave roofed or covered areas, being able to be spotted easily from an UAV. In the infrared spectrum, it is possible to detect thermal signatures of animals in medium, broken and sparse canopy [46]. Infrared thermal imagery is therefore implementable in more various settings than the visible spectrum. Automatic detection of the dogs using identification and counting algorithms or deep learning could be used to identify and count dogs on the UAV’s pictures in both visible and infrared spectrum. Algorithms have already been developed for red deer, roe deer, wild boars [34], bison, elks, wolves [35], koalas, kangaroos [31] and cows [47]. Deep learning has been used for automatically identifying, counting and describing wild animals in camera-trap images [48]. Although, the study implemented by Chrétien et al was not completely successful for the identification of wolves in UAV pictures using a multicriteria object-based image analysis [35], the utilization of artificial intelligence in the study conducted by Gonzalez et al showed good results for small size animals like koalas [31]. Thanks to this progress, we are confident that such methods can be developed for dogs. This would fasten the pictures’ analysis and considerably lower the workload, which is one of the drawback of using UAV in animal management and research [31].

The third possibility is to combine UAV flight and capture-recapture methods. Such a technique accounts for the fact that it is impossible to detect all the dogs during the UAV flight. Dogs can be marked using a technique easily visible by the UAV, for example red paint on the back of the dogs, and captured again during the UAV flights. The Bayesian model developed within this study demonstrated its performance well and could be used in combination with the UAV to obtain closer estimates of dog population sizes.

The flight altitude of the UAV should be low enough to allow the identification of dogs. The mountainous landscape in La Romana prevented the UAV from flying below 100 meters. UAV’s flight should be performed in flat areas, without hills and other obstacles (e.g. antennas) to facilitate low-altitude flights. Using flight planning software allowing for terrain following (e.g. UgCS www.ugcs.com or Map Pilot www.mapsmadeeasy.com) could be an option for uneven areas. These software allow the UAV to fly at a constant level about the ground, regardless of the topography, instead of flying at a constant altitude. According to Chrétien et al, wolves could be confused with rocks for flights at 60 meters [35]. Optimal flight altitudes should be identified for the respective species and environment before the use of UAVs.

The reasons for poor quality of the pictures usually were the weather conditions. During the UAV’s flight, the weather needs to be clear without wind, which causes UAV flight’s instability and blurred pictures.

An important point to take into consideration is the prevalent gray area regarding country regulations when employing UAVs in research related to public health, especially when flying over private and public areas at low altitudes, as it elicits risks for breach of privacy and public safety. In applications such as this one, ethical approval has to be granted by ERBs who might demand to follow compliance with detailed protocols of human subjects’ research, even if no direct intervention with people (such as interviews) is foreseen. This can be a cumbersome process if no precedents are established in the country or study area, increasing time and resources needed to comply with varied interpretations of an appropriate ethical standard. However, to ensure a best practice approach in this emerging field of application, it is recommended to be overly cautious in securing local communities’ formal approval of the research intervention, being mindful to sociocultural expectations, traditional governance, and consuetudinary law.

### Method 2: Bayesian statistical model based on capture-recapture data

The capture-recapture technique and analysis using Bayesian statistical model estimated the size of FRDD populations in a satisfactory way in this study. The presence of puppies can explain the difference between the estimate given by this method and the dog census. Capture-recapture methods already demonstrated efficiency in the literature. It is simple method to provide population size estimates [17] and they are useful to understand population dynamics of FRDD [20]. The use of a Bayesian statistical model is able to capture heterogeneity in the detection probabilities due to confinement of owned dogs and presence of ownerless dogs.

To assess the impact of the prior values (i.e. confinement probability, ownerless to owned dog ratio and number of owned dogs living in the study site) on the three outcomes (i.e. number of owned FRDD, number of ownerless dogs and total number of FRDD), we increased the range of the confinement probability uniform distribution, the expected value of the ownerless to owned dogs ratio (log normal distribution) and the range of the number of owned dog uniform distribution. The maximum value of confinement probability was increased to 0.5 in La Romana and Sabaneta and to 0.75 in Poptún. The three outcomes were impacted of less than 1%. A gamma distribution, allowing a higher ownerless to owned dog ratio than the log normal distribution used previously in the model, was used to assess the impact of this prior. The choice of the prior distribution had an impact on the estimated number of owned FRDD and ownerless dogs but had an impact of maximum 3% on the total number of FRDD (Table 3 – Supporting information). This limitation of relying on prior values for the proportion of owned to ownerless dogs is not critical if the aim is to estimate the total size of FRDD population. This model should not be used to calculate the size of owned dog population in a region where ownerless dogs are present and there are not reliable prior information on ownerless dog population. The increase of the maximum value of owned dog distribution had an impact of less than 1% on the outcomes. This method performed well in counting free-roaming dogs, especially when ownerless dogs are present and prior information is available. However the drawback of not counting puppies remains.

### Method 3: the human:dog ratio

The human:dog ratio method is currently used by the health authorities in Guatemala to estimate the number of vaccines needed for national rabies vaccination campaigns. While this method produced relatively close results for La Romana, it highly underestimated the population size in Sabaneta, taking the dog census data as a reference. The use of the dog and human census data quantified by “One Health Poptún” project to compute the human:dog ratio results in a value 4.4:1 in La Romana and 2.7:1 in Sabaneta. Although Sabaneta and La Romana are both in a rural setting and located only 30km apart, the human:dog ratio varies strongly. This highlights that an overall country human:dog ratio is not the best way to estimate size of FRDD populations. These findings are consolidated by the results of the dog census performed in Todos Santos Cuchumatán neighbourhoods which results in a human:owned dog ratio varying from 4:1 to 12.3:1 [27] and the findings of the studies currently ongoing at the UVG (unpublished data). It would be interesting to assess if the human:dog ratio is dependent on factors such as socioeconomic or environmental factors. Studies performed in Africa indicated that religion and the need for guarding property and protecting livestock could be part of those factors [49,50].

## Conclusion

This is the first study published on the application of UAV to estimate population size of free roaming domestic dogs and it could be shown that the technique worked to detect and count dog in the pictures. However, the sensitivity of the technique has to be improved, for example via a wise choice of the timing and the study area, the utilization of infrared thermal imagery, the utilization of automatic detection of the dogs or the combination of UAV and capture recapture methods. This study may be useful to give directions on how to design efficient application of UAV for this purpose. It also highlights that the aim for the dog populations size estimation influence the method to be selected.

## Acknowledgments

We thank Pablo Ax, Claudia Caal, Pablo Chub, Julio Eufragio, Dione Méndez, Milton Oliva, Jorge Paniagua, German Rodas, and Alexis Tut for their help and excellent work on the field; Wendy Hernández, Ramón Medrano and Adan Real for reviewing the drone pictures; Sonja Hartnack and Beatriz Vidondo for their feedback on the Bayesian statistical model; Jakob Zinsstag for giving us the opportunity to use the “One Health Poptún” project setting to implement this study and all the dog owner’s for their participation.

## Supporting information

**S1 Fig. Transects lines in La Romana, Petén, Guatemala**. The green and the red dots are the starting and ending points, respectively.

**S2 Fig. Transects lines in Sabaneta, Petén, Guatemala**. The green and the red dots are the starting and ending points, respectively.

**S3 Fig. Transects lines in Poptun, Petén, Guatemala**. The green and the red dots are the starting and ending points, respectively, the red square depicts the 1km^2^ study site.

**S4 Fig. Ratio of estimated number of ownerless to owned dogs in Poptún, Guatemala.**

**S1 Table. Confinement practices of dog’s owners in Petén department, Guatemala**.

**S2 Table. Number of marked and unmarked dogs recaptured during the ground transects**.

**S3 Table. Impact of the prior ownerless to owned dogs ratio distribution on the model outcomes.**

**S1 File. Script of the Bayesian statistical model implemented on OpenBUGS**.

## References

1. Creel S, Spong G, Sands JL, Rotella J, Zeigle J, Joe L, et al. Population size estimation in Yellowstone wolves with error-prone noninvasive microsatellite genotypes. Mol Ecol. 2003;12(7):2003–9.

2. Katzner TE, Ivy JAR, Bragin EA, Milner-Gulland EJ, Dewoody JA. Conservation implications of inaccurate estimation of cryptic population size. Anim Conserv. 2011;14(4):328–32.

3. Stallknecht DE. Impediments to wildlife disease surveillance, research, and diagnostics. Curr Top Microbiol Immunol. 2007;315:445–61.

4. Yoccoz NG, Nichols JD, Boulinier T. Monitoring of biological diversity in space and time. Trends Ecol Evol. 2001;16(8):446–53.

5. Williams R, Hedley SL, Branch TA, Bravington M V., Zerbini AN, Findlay KP. Chilean Blue Whales as a Case Study to Illustrate Methods to Estimate Abundance and Evaluate Conservation Status of Rare Species. Conserv Biol. 2011;25(3):526–35.

6. Gerber BD, Ivan JS, Burnham KP. Estimating the abundance of rare and elusive carnivores from photographic-sampling data when the population size is very small. Popul Ecol. 2014;56(3):463–70.

7. Tenzin T, Ahmed R, Debnath NC, Ahmed G, Yamage M. Free-Roaming Dog Population Estimation and Status of the Dog Population Management and Rabies Control Program in Dhaka City, Bangladesh. PLoS Negl Trop Dis [Internet]. 2015;9(5):1–14. Available from: http://dx.doi.org/10.1371/journal.pntd.0003784

8. Byrne AW, O’Keeffe J, Green S, Sleeman DP, Corner LAL, Gormley E, et al. Population Estimation and Trappability of the European Badger (Meles meles): Implications for Tuberculosis Management. PLoS One. 2012;7(12).

9. Kayali U, Mindekem R, Hutton G, Ndoutamia AG, Zinsstag J. Cost-description of a pilot parenteral vaccination campaign against rabies in dogs in N’Djaména, Chad. Trop Med Int Heal. 2006;11(7):1058–65.

10. Murphy SM, Wilckens DT, Augustine BC, Peyton MA, Harper GC. Improving estimation of puma (Puma concolor) population density: clustered camera-trapping, telemetry data, and generalized spatial mark-resight models. Sci Rep [Internet]. 2019;9(1):1–13. Available from: http://dx.doi.org/10.1038/s41598-019-40926-7

11. Downes MJ, Dean RS, Stavisky JH, Adams VJ, Grindlay DJC, Brennan ML. Methods used to estimate the size of the owned cat and dog population: A systematic review. BMC Vet Res. 2013;9.

12. Alves MCGP, Matos MR de, Reichmann M de L, Dominguez MH. Estimation of the dog and cat population in the State of São Paulo. Rev Saude Publica [Internet]. 2005;39(6):891–7. Available from: http://www.ncbi.nlm.nih.gov/pubmed/16341397

13. Belo VS, Werneck GL, Da Silva ES, Barbosa DS, Struchiner CJ. Population estimation methods for free-ranging dogs: A systematic review. PLoS One. 2015;10(12):1–15.

14. Childs JE, Robinson LE, Sadek R, Madden A, Miranda ME, Miranda NL. Density estimates of rural dog populations and an assessment of marking methods during a rabies vaccination campaign in the Philippines. Prev Vet Med. 1998;33(1–4):207–18.

15. Gsell AS, Knobel DL, Sarah C, Kazwala RR, Vounatsou P, Zinsstag J. Domestic dog demographic structure and dynamics relevant to rabies control planning in urban areas in Africa: the case of Iringa, Tanzania. BMC Vet Res. 2012;8(1):236.

16. Tiwari HK, Vanak AT, O’Dea M, Gogoi-Tiwari J, Robertson ID. A Comparative Study of Enumeration Techniques for Free-Roaming Dogs in Rural Baramati, District Pune, India. Front Vet Sci. 2018;5(May):1–12.

17. Hiby LR, Reece JF, Wright R, Jaisinghani R, Singh B, Hiby EF. A mark-resight survey method to estimate the roaming dog population in three cities in Rajasthan, India. BMC Vet Res [Internet]. 2011;7(1):46. Available from: http://www.biomedcentral.com/1746-6148/7/46

18. World Health Organization. WHO expert consultation on rabies. World Health Organization - Technical Report Series. 2018.

19. World Organization for Animal Health (OIE). Guidelines on stray dog population control. OIE Terr Anim Heal Stand Comm [Internet]. 2009;Chap 7.7.:313–621. Available from: https://www.oie.int/doc/ged/D9926.PDF

20. Belo VS, Struchiner CJ, Werneck GL, Teixeira Neto RG, Tonelli GB, De Carvalho CG, et al. Abundance, survival, recruitment and effectiveness of sterilization of free-roaming dogs: A capture and recapture study in Brazil. PLoS One. 2017;12(11):1–19.

21. Hossain M, Ahmed K, Marma ASP, Hossain S, Ali MA, Shamsuzzaman AKM, et al. A survey of the dog population in rural Bangladesh. Prev Vet Med [Internet]. 2013;111(1–2):134–8. Available from: http://dx.doi.org/10.1016/j.prevetmed.2013.03.008

22. Aiyedun J, Olugasa B. Use of aerial photograph to enhance dog population census in Ilorin, Nigeria. Sokoto J Vet Sci. 2012;10(1):22–7.

23. Buckland ST, Anderson DR, Burnham KP, Laake JL. Distance Sampling. In: Encyclopedia of Biostatistics. Second edi. 2005.

24. Kayali U, Mindekem R, Yémadji N, Vounatsou P, Kaninga Y, Ndoutamia AG, et al. Coverage of pilot parenteral vaccination campaign against canine rabies in N’Djaména, Chad. Bull World Health Organ [Internet]. 2003;81(10):739–44. Available from: http://www.ncbi.nlm.nih.gov/pubmed/14758434%0Ahttp://www.pubmedcentral.nih.gov/articlerender.fcgi?artid=PMC2572337

25. Dürr S, Ward MP. Development of a novel rabies simulation model for application in a non-endemic environment. PLoS Negl Trop Dis. 2015;9(6).

26. Ministerio de Salud Pública y asistencia Social de Guatemala MSPAS. Situación Epidemiológica de Rabia Antecedentes en Guatemala. In 2017.

27. Pulczer AS, Jones-Bitton A, Waltner-Toews D, Dewey CE. Owned dog demography in Todos Santos Cuchumatán, Guatemala. Prev Vet Med [Internet]. 2013;108(2–3):209–17. Available from: http://dx.doi.org/10.1016/j.prevetmed.2012.07.012

28. Faillace De Leon R. La campaña antirrabica en Guatemala. Bol la Of Sanit Panam. 1959;(April):356–60.

29. Hodgson JC, Baylis SM, Mott R, Herrod A, Clarke RH. Precision wildlife monitoring using unmanned aerial vehicles. Sci Rep [Internet]. 2016;6(March):1–7. Available from: http://dx.doi.org/10.1038/srep22574

30. Díaz-Delgado R, Hurford C, Lucas R. Introducing the Book “The Roles of Remote Sensing in Nature Conservation.” The Roles of Remote Sensing in Nature Conservation. 2017. 3–10 p.

31. Gonzalez LF, Montes GA, Puig E, Johnson S, Mengersen K, Gaston KJ. Unmanned aerial vehicles (UAVs) and artificial intelligence revolutionizing wildlife monitoring and conservation. Sensors (Switzerland). 2016;16(1).

32. Zmarz A, Korczak-Abshire M, Storvold R, Rodzewicz M, Kȩdzierska I. Indicator species population monitoring in Antarctica with UAV. Int Arch Photogramm Remote Sens Spat Inf Sci - ISPRS Arch. 2015;40(1W4):189–93.

33. Seymour AC, Dale J, Hammill M, Halpin PN, Johnston DW. Automated detection and enumeration of marine wildlife using unmanned aircraft systems (UAS) and thermal imagery. Sci Rep [Internet]. 2017;7(March):1–10. Available from: http://dx.doi.org/10.1038/srep45127

34. Witczuk J, Pagacz S, Zmarz A, Cypel M. Exploring the feasibility of unmanned aerial vehicles and thermal imaging for ungulate surveys in forests - preliminary results. Int J Remote Sens [Internet]. 2018;39(15–16):5504–21. Available from: https://doi.org/10.1080/01431161.2017.1390621

35. Chrétien LP, Théau J, Ménard P. Wildlife multispecies remote sensing using visible and thermal infrared imagery acquired from an unmanned aerial vehicle (UAV). Int Arch Photogramm Remote Sens Spat Inf Sci - ISPRS Arch. 2015;40(1W4):241–8.

36. McClelland GTW, Bond AL, Sardana A, Glass T. Unmanned aerial vehicles for population estimates. Mar Ornithol. 2016;44:215–20.

37. Comerford SC. Medicinal plants of two Mayan healers from San Andrés, Petén, Guatemala. Econ Bot. 1996;50(3):327–36.

38. Beach T. Soil catenas tropical deforestation, and ancient and contemporary soil erosion in the petén, guatemala. Phys Geogr [Internet]. 1998;19(5):378–405. Available from: 10.1080/02723646.1998.10642657

39. Torres D. Trabajo de tesis de Ingeniería Ambiental: Análisis de las causas y agentes de la deforestación en sabaneta, Poptún, Petén, para el diseño de un plan de manejo forestal a nivel comunitario. Universidad Rafael Landívar: Guatemala; 2019.

40. Instituto Nacional de Estadística –INE. Resultados del Censo Nacional de Población 2018. República de Guatemala: Guatemala. [Internet]. Available from: https://www.censopoblacion.gt

41. Stöcker C, Bennett R, Nex F, Gerke M, Zevenbergen J. Review of the current state of UAV regulations. Remote Sens. 2017;9(5):33–5.

42. World Medical Association (WMA). Declaration of Helsinki-Ethical Principles for Medical Research Involving Human Subjects [Internet]. 59th WMA General Assembly, Seoul, Korea; 2008. Available from: https://www.wma.net/policies-post/wma-declaration-of-helsinki-ethical-principles-for-medical-research-involving-human-subjects/2013/

43. Cosenza C, Eudey L, Kerr J, Trumbo B. A Review of Methods for Point and Interval Estimation of Population Size in Capture-Recapture Studies. 2014;2(1).

44. Van Hauwermeiren M, Vose D. A compendium of distributions [ebook]. Vose Software, Ghent, Belgium [Internet]. 2009. 1–173 p. Available from: http://www.vosesoftware.com

45. Van Bommel L, Johnson CN. Where do livestock guardian dogs go? Movement patterns of free-ranging Maremma sheepdogs. PLoS One. 2014;9(10):1–12.

46. Witczuk J, Pagacz S, Zmarz A, Cypel M. Exploring the feasibility of unmanned aerial vehicles and thermal imaging for ungulate surveys in forests - preliminary results. Int J Remote Sens. 2018;39(15–16):5504–21.

47. Longmore SN, Collins RP, Pfeifer S, Fox SE, Mulero-Pázmány M, Bezombes F, et al. Adapting astronomical source detection software to help detect animals in thermal images obtained by unmanned aerial systems. Int J Remote Sens [Internet]. 2017;38(8–10):2623–38. Available from: http://dx.doi.org/10.1080/01431161.2017.1280639

48. Norouzzadeh MS, Nguyen A, Kosmala M, Swanson A, Palmer M, Packer C, et al. Automatically identifying, counting, and describing wild animals in camera-trap images with deep learning. 2017;115(25). Available from: http://arxiv.org/abs/1703.05830

49. Mbilo C, Léchenne M, Hattendorf J, Madjadinan S, Anyiam F, Zinsstag J. Rabies awareness and dog ownership among rural northern and southern Chadian communities—Analysis of a community-based, cross-sectional household survey. Acta Trop [Internet]. 2017;175:100–11. Available from: http://dx.doi.org/10.1016/j.actatropica.2016.06.003

50. Ratsitorahina M, Rasambainarivo JH, Raharimanana S, Rakotonandrasana H, Andriamiarisoa MP, Rakalomanana FA, et al. Dog ecology and demography in Antananarivo, 2007. BMC Vet Res. 2009;5:1–7.

